# Neurostructural correlates of real-life social drinking risk in alcohol use disorder

**DOI:** 10.64898/2026.10.13.758538

**Authors:** Ali Ghadami, Edda Bilek, Iris Reinhard, Abhijit Sreepada, Marvin Ganz, Jamila Andoh, Justin Boehmer, Kanya Meyer-Degering, Dominic Reichert, Henrik Walter, Bernd Lenz, Patrick Bach, Falk Kiefer, Markus Reichert, Ulrich Ebner-Priemer, Andreas Meyer-Lindenberg, Christine Heim, Heike Tost

**Affiliations:** Department of Psychiatry and Psychotherapy, Central Institute of Mental Health, University of Heidelberg, Medical Faculty Mannheim, Mannheim, Baden-Württemberg, Germany; German Center for Mental Health (DZPG), Partner Site Mannheim-Heidelberg-Ulm, Germany; Core Facility Biostatistics, Central Institute of Mental Health, University of Heidelberg, Medical Faculty Mannheim, Mannheim, Baden-Württemberg, Germany; Charité-Universitätsmedizin Berlin, Corporate Member of Freie Universität Berlin and Humboldt-Universität zu Berlin, Institute of Medical Psychology, Berlin, Germany; Charité-Universitätsmedizin Berlin, Corporate Member of Freie Universität Berlin and Humboldt-Universität zu Berlin, Department of Psychiatry and Psychotherapy (CCM), Berlin, Germany; German Center for Mental Health (DZPG), Partner Site Berlin-Potsdam, Germany; Department of Addictive Behavior and Addiction Medicine, Central Institute of Mental Health, Medical Faculty Mannheim, Heidelberg University, Mannheim, Germany; Department of Sport and Exercise Science, Faculty of Natural and Life Sciences, University of Salzburg, Salzburg, Austria; Mental mHealth Lab, Department of Applied Psychology, Institute of Sports and Sports Science, Karlsruhe Institute of Technology, Karlsruhe, Germany; Charité-Universitätsmedizin Berlin, Corporate Member of Freie Universität Berlin and Humboldt-Universität zu Berlin, NeuroCure Cluster of Excellence, Berlin, Germany

## Abstract

The social context of alcohol consumption is increasingly recognized as an important factor in the development of alcohol use disorder (AUD). However, individuals with AUD may differ in how strongly alcohol use by people around them is associated with their own drinking, and the neurostructural basis of this variation remains unclear. Here we studied 83 individuals with AUD (53 men, 30 women; mean [SD] age, 37.20 [12.42] years; median DSM-5 AUD criteria, 4) using six weeks of ecological momentary assessment and structural magnetic resonance imaging (n = 80). Across 5,221 non-solitary observations, greater exposure to surrounding drinkers was associated with increased odds of momentary drinking (odds ratio, 4.23; 95% CI, 3.60-4.97), with significant between-person variation that we quantified as social drinking risk (SDR). Higher SDR was associated with lower gray matter volume in the pregenual anterior cingulate cortex (T = 5.40; P < .001) and weaker pregenual anterior cingulate cortex–ventral striatum structural covariance (T = 3.15; P = .032). These findings support SDR as an ecologically grounded phenotype linking daily-life social-context sensitivity to cingulostriatal structure, with potential relevance for mechanistic and interventional research in AUD.

## Introduction

Alcohol use commonly occurs in social settings, and social context contributes to the development and maintenance of alcohol use disorder (AUD) ^1,2^. Social drinking is linked to enhanced positive emotions and social experiences, and drinking with others is associated with greater alcohol consumption and stronger rewarding effects than drinking alone ^2,3^. These associations are not uniform across individuals ^1,4^. Group-average findings may obscure individuals whose drinking is particularly responsive to surrounding alcohol use. Such heterogeneity is difficult to capture using retrospective reports or laboratory paradigms. Ecological momentary assessment (EMA) is therefore well suited to quantify person-specific sensitivity to daily-life social drinking contexts ^3,4^.

Daily-life social-context sensitivity may relate to brain systems that integrate social, affective, and contextual information with reward processing ^5-7^. The pregenual anterior cingulate cortex (pgACC) is a key node in this circuitry ^7^ and has been proposed as an integrative locus through which social-environmental influences contribute to psychiatric risk ^8^. Our previous work showed that greater affective benefit from real-life social contact is associated with higher pgACC gray matter volume (GMV) ^9^, whereas AUD is reportedly associated with lower cingulate GMV ^10^. Anatomical and functional imaging studies further link the pgACC and ventral striatum (VS) within circuitry implicated in reward processing and alcohol cue reactivity ^6,11,12^. However, whether individual differences in daily-life social-context sensitivity relate to the pgACC and its interaction with the VS in AUD remains unclear.

We studied individuals with AUD using 6 weeks of smartphone-based EMA and structural magnetic resonance imaging (MRI). We derived a social drinking risk (SDR) index reflecting how strongly each participant’s momentary drinking likelihood increased with exposure to surrounding drinkers in real life. Based on our prior work ^9^ and related findings, we hypothesized that higher SDR would be associated with lower pgACC GMV and weaker pgACC-VS structural covariance.

## Materials and methods

### Study Design and Participants

We recruited 83 individuals (53 men, 30 women; mean [SD] age, 37.20 [12.42] years) meeting 2 to 8 DSM-5 AUD criteria ^13^ (mean [SD], 4.02 [1.59]; median, 4) between March 2021 and March 2023. Participants underwent structural MRI followed by 6 weeks of smartphone-based EMA between 08:00 and 22:00 (4-8 prompts/day). T1-weighted 3D MPRAGE images were acquired on a 3-T Siemens Prismafit scanner at 1.0-mm isotropic resolution (details in Supplementary information). Prompts assessed momentary drinking (yes/no) and, in non-solitary contexts, exposure to surrounding drinkers (none, a few, about half, majority, all; coded 1-5; Figure 1A). Solitary observations were excluded because the surrounding-drinkers item was inapplicable. All participants provided written informed consent and received monetary compensation.

**FIGURE 1.**
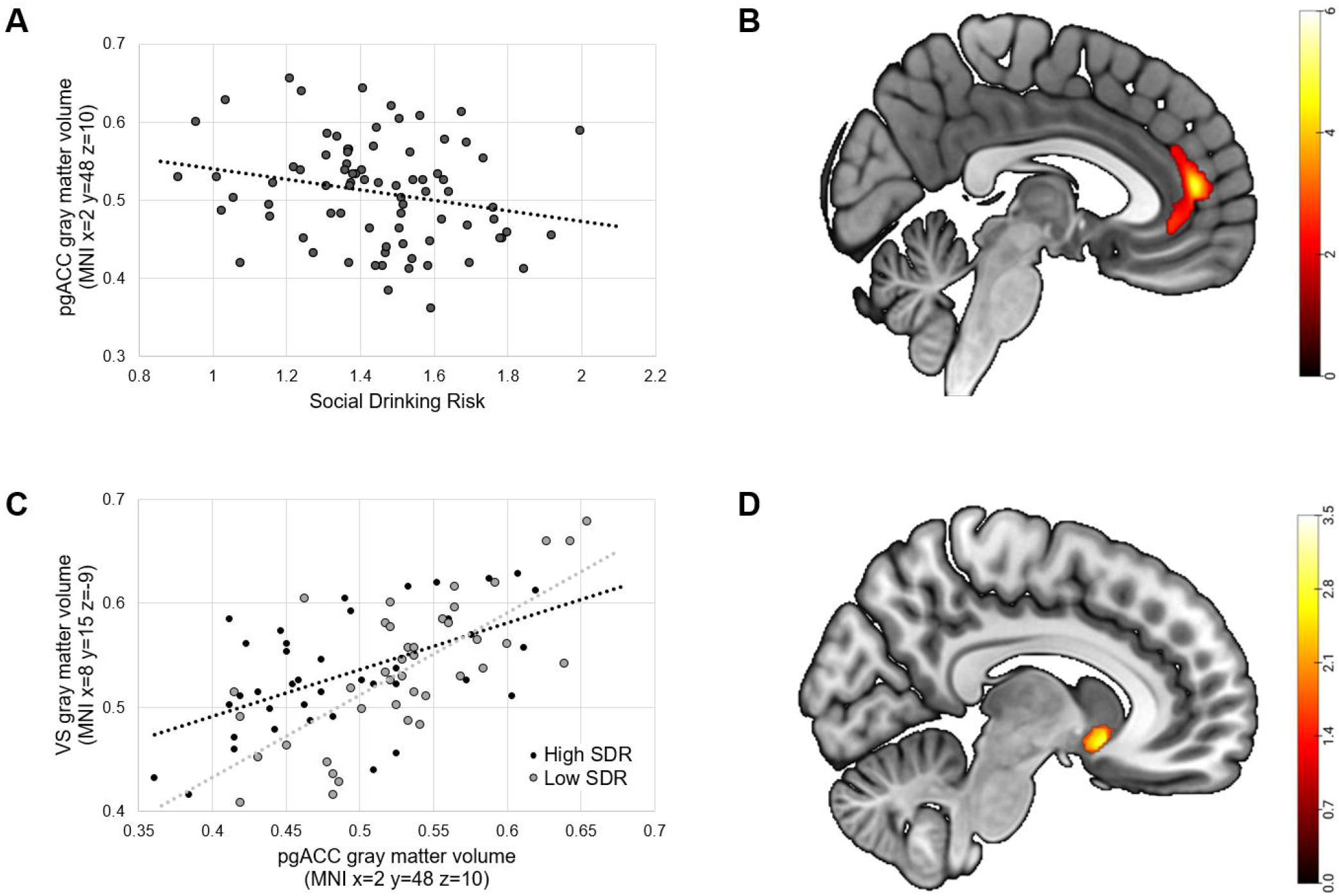
Title: Exposure to surrounding drinkers and predicted momentary drinking. **A**. Illustration of EMA items on a representative smartphone (shown on a single screen for illustrative purposes). **B**. Predicted probability of momentary drinking (outcome variable; y-axis) according to exposure to surrounding drinkers (predictor; x-axis). Predicted probabilities at each surrounding-drinking category were calculated for each participant. Gray lines show participant-specific predictions and the black line shows the fixed-effects prediction averaged across participants at each exposure level. The corresponding predictions on the log-odds scale are presented in Figure S1.

### Statistical Analysis

We used a generalized linear mixed model (GLMM) with a logit link in SAS to test the association between exposure to surrounding drinkers (coded 1-5; continuous predictor) and momentary drinking (main outcome). The model included participant-specific random intercepts and slopes for exposure to surrounding drinkers, adjusted for age, sex, and time of day (Equation S1). We extracted participant-specific slope values (fixed-effect estimate plus participant-specific random slope estimate) for the association between exposure and momentary drinking likelihood. These slopes defined SDR, which quantified the extent to which a participant’s drinking likelihood increased with higher exposure to surrounding drinkers. SDR was used as the predictor in subsequent MRI analyses. Figure 1B shows GLMM-predicted probabilities, evaluated at each participant’s sex and age and a fixed time of day (18:00); corresponding log-odds predictions are shown in Figure S1. Voxel-based morphometry (VBM) tested the association between SDR and pgACC GMV, and structural covariance analysis tested whether SDR moderated the association between peak-voxel pgACC GMV and VS GMV; both models adjusted for age, sex, and total GMV. Statistical inference used family-wise error (FWE) small-volume correction within prespecified pgACC and VS masks, respectively.

## Results

The EMA analysis included 12,774 observations (median [IQR], 163 [108-196] per participant). The GLMM included 5,221 non-solitary observations (median [IQR], 51 [28-87] per participant). Higher surrounding-drinkers exposure was associated with higher odds of momentary drinking (odds ratio [OR], 4.23; 95% CI, 3.60-4.97; *P* < .001). This association varied significantly across participants (random-slope variance, 0.14; 95% Wald confidence bounds, 0.07-0.37; P = .005), indicating person-specific differences in social-context sensitivity (Table S1). To illustrate this association on the probability scale, fixed-effects predictions averaged across participants ranged from 6% when no surrounding people were drinking to 94% when all were drinking (Figure 1B). SDR was not associated with alcohol use severity, social network measures, or other baseline variables after false discovery rate correction (Spearman ρ range, −0.13 to 0.27; adjusted *P* ≥ .11; Table S2).

In the MRI subsample (n = 80), higher SDR was associated with lower pgACC GMV (MNI peak, [2, 48, 10]; T = 5.40; *P* < .001, FWE small-volume corrected within the prespecified pgACC mask; Figure 2A and B). A significant pgACC-by-SDR interaction in the VS indicated weaker pgACC-VS structural covariance at higher SDR (MNI peak, [8, 15, −9]; T = 3.15; *P* = .032, FWE small-volume corrected within the prespecified VS mask; Figure 2C and D).

**FIGURE 2.**
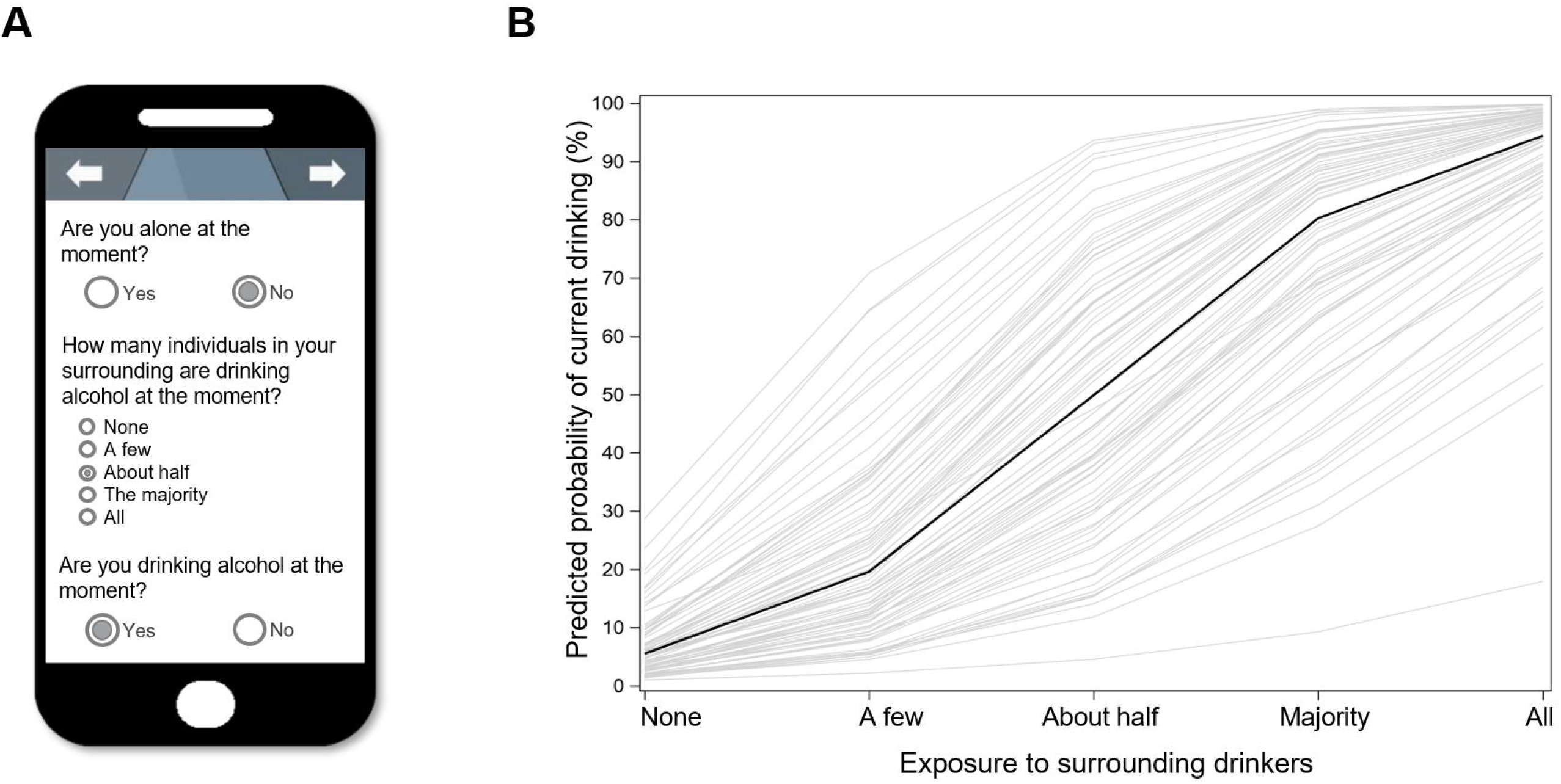
Title: Structural brain associations with social drinking risk. **A**. Illustrative association between SDR and peak-voxel pgACC GMV. **B**. Sagittal view at MNI x = 3 mm, selected to display the pgACC cluster clearly; the statistical peak was [2, 48, 10]. The map is displayed at P < .05, uncorrected, for visualization. **C**. Illustrative association between peak-voxel pgACC and VS GMV at the structural covariance FWE-SVC peak ([8, 15, −9]), split by median SDR (high vs. low; display only). **D**. Sagittal view at MNI x = 8 mm, centered on the VS, showing the pgACC-by-SDR interaction, displayed at P < .05, uncorrected for visualization. The color bars represent the t-statistic. Statistical inference used family-wise error small-volume correction within the predefined pgACC and VS masks.

## Discussion

Greater exposure to surrounding drinkers was associated with higher odds of momentary drinking in AUD, with significant between-person variation quantified as SDR. These findings extend evidence linking cue-rich social contexts to alcohol reward and drinking ^1–4^ by demonstrating substantial individual differences in susceptibility to surrounding alcohol use in real-life settings. SDR therefore provides a novel, ecologically grounded, person-specific measure of social-context sensitivity. The absence of significant associations with alcohol use severity, social support, or social-network measures further suggests that SDR may capture a complementary dimension of context-dependent drinking risk. Future studies should examine whether social anxiety, drinking motives, perceived norms, or sensitivity to alcohol’s social rewards explain variation in SDR.

The association of higher SDR with lower pgACC GMV extends evidence of cingulate alterations in AUD ^10^ and prior work linking pgACC structure to affective responses in real-life social contexts ^9^, suggesting that variation in this socially and affectively relevant region ^7^ may contribute to individual drinking risk in cue-rich social environments. Higher SDR was also associated with weaker pgACC-VS structural covariance, consistent with cingulostriatal involvement in alcohol cue and reward processing ^6,7,11,12^. Because structural covariance reflects coordinated interindividual variation in regional morphology ^14^, lower pgACC-VS covariance at higher SDR may indicate altered structural organization of pgACC-VS circuitry. One possible functional interpretation is reduced top-down control relevant to alcohol cue processing and motivated behavior ^6,7,12^, which could permit disinhibited reward-expectancy signaling and contribute to increased alcohol consumption. Together, these findings suggest that cingulostriatal structural variation is relevant to socially cued drinking in AUD ^6,8,15^.

The potential clinical value of SDR may lie in distinguishing individuals with AUD whose drinking is especially responsive to social alcohol cues from those whose risk is driven by other contexts. If prospectively validated, SDR could inform context-focused skills training and just-in-time adaptive support during high-risk social situations. Longitudinal and interventional studies should test whether SDR predicts relapse and whether SDR-informed interventions improve outcomes.

Limitations include the observational design, which precludes causal inference regarding both the exposure-drinking association and whether neural differences precede or follow context-sensitive drinking. Structural covariance reflects coordinated interindividual variation, not anatomical or functional connectivity, and requires validation using functional and structural connectivity measures. Despite these limitations, SDR is a candidate ecologically grounded phenotype linking daily-life social drinking context to cingulostriatal structure in AUD.

## Ethics approval and consent to participate

The study protocol was approved by the Medical Ethics Committee II of the Medical Faculty Mannheim, Heidelberg University (reference no. 2018-621N-MA). All methods were performed in accordance with the relevant guidelines and regulations, including the Declaration of Helsinki. All participants provided written informed consent and received monetary compensation.

## Acknowledgments

We thank all individuals who supported our work by participating in the study. We thank Mirjam Melzer, Beate Hoechemer, and our research assistants for their valuable help with data acquisition.

## Author Contributions

A.G., A.S., M.G., J.B., and K.M.-D. acquired the data. A.G., I.R., D.R., and M.R. analyzed the data. A.G. drafted the manuscript. H.T. supervised the study and contributed to interpreting the results and writing the manuscript. H.T., U.W.E.-P., and C.H. designed the study and acquired funding. A.G., A.S., I.R., M.G., J.B., K.M.-D., H.W., J.A., B.L., P.B., F.K., E.B., M.R., U.W.E.-P., A.M.-L., C.H., and H.T. contributed to data interpretation and critically revised the manuscript. All authors reviewed and approved the final manuscript.

## References

1 Fairbairn, C. E. & Kang, D. Social Drinking and Addiction: A Social-Cognitive Model for Understanding Alcohol Use Disorder Risk. Curr Dir Psychol Sci (2025). 10.1177/09637214251318272

2 Creswell, K. G. Drinking Together and Drinking Alone: A Social-Contextual Framework for Examining Risk for Alcohol Use Disorder. Curr Dir Psychol Sci 30, 19–25 (2021). 10.1177/0963721420969406

3 Fischer, A. M., King, A. C., Cao, D. & Fridberg, D. J. Drinking context, alcohol use, and subjective responses during binge drinking episodes measured by high-resolution ecological momentary assessment (HR-EMA). Psychol Addict Behav 37, 258–266 (2023). 10.1037/adb0000887

4 Acuff, S. F., Padovano, H. T., Carpenter, R. W., Emery, N. N. & Miranda, R., Jr. Effects of social drinking context on subjective effects, affect, and next-day appraisals in the natural environment. Alcohol Clin Exp Res (Hoboken) 48, 755–765 (2024). 10.1111/acer.15286

5 Apps, M. A., Rushworth, M. F. & Chang, S. W. The Anterior Cingulate Gyrus and Social Cognition: Tracking the Motivation of Others. Neuron 90, 692–707 (2016). 10.1016/j.neuron.2016.04.018

6 Haber, S. N. & Knutson, B. The reward circuit: linking primate anatomy and human imaging. Neuropsychopharmacology 35, 4–26 (2010). 10.1038/npp.2009.129

7 Palomero-Gallagher, N., Hoffstaedter, F., Mohlberg, H., Eickhoff, S. B., Amunts, K. & Zilles, K. Human Pregenual Anterior Cingulate Cortex: Structural, Functional, and Connectional Heterogeneity. Cereb Cortex 29, 2552–2574 (2019). 10.1093/cercor/bhy124

8 Meyer-Lindenberg, A. & Tost, H. Neural mechanisms of social risk for psychiatric disorders. Nat Neurosci 15, 663–668 (2012). 10.1038/nn.3083

9 Gan, G., Ma, R., Reichert, M., Giurgiu, M., Ebner-Priemer, U. W., Meyer-Lindenberg, A. et al. Neural Correlates of Affective Benefit From Real-life Social Contact and Implications for Psychiatric Resilience. JAMA Psychiatry 78, 790–792 (2021). 10.1001/jamapsychiatry.2021.0560

10 Li, L., Yu, H., Liu, Y., Meng, Y. J., Li, X. J., Zhang, C. et al. Lower regional grey matter in alcohol use disorders: evidence from a voxel-based meta-analysis. BMC Psychiatry 21, 247 (2021). 10.1186/s12888-021-03244-9

11 Schacht, J. P., Anton, R. F. & Myrick, H. Functional neuroimaging studies of alcohol cue reactivity: a quantitative meta-analysis and systematic review. Addict Biol 18, 121–133 (2013). 10.1111/j.1369-1600.2012.00464.x

12 Zeng, J., Yu, S., Cao, H., Su, Y., Dong, Z. & Yang, X. Neurobiological correlates of cue-reactivity in alcohol-use disorders: A voxel-wise meta-analysis of fMRI studies. Neurosci Biobehav Rev 128, 294–310 (2021). 10.1016/j.neubiorev.2021.06.031

13 American Psychiatric Association. Diagnostic and Statistical Manual of Mental Disorders. Fifth edition. edn, (American Psychiatric Association, 2013).

14 Mechelli, A., Friston, K. J., Frackowiak, R. S. & Price, C. J. Structural covariance in the human cortex. J Neurosci 25, 8303–8310 (2005). 10.1523/JNEUROSCI.0357-05.2005

15 Beckmann, M., Johansen-Berg, H. & Rushworth, M. F. Connectivity-based parcellation of human cingulate cortex and its relation to functional specialization. J Neurosci 29, 1175–1190 (2009). 10.1523/JNEUROSCI.3328-08.2009

