## Supplemental information for "Neurostructural correlates of real-life social drinking risk in alcohol use disorder"

### 1. Supplementary Methods

#### 1.1. Participants and study procedure

Participants were recruited from the main cohort of the TRR 265 addiction research consortium. The TRR 265 is a multidisciplinary research consortium dedicated to neurobiological, psychological, and social mechanisms contributing to substance use disorders (https://www.trr265.org/en)[1]. Eligible participants met 2 to 8 Alcohol Use Disorder (AUD) criteria (mean [SD], 4.02 [1.59]; median, 4) according to the Diagnostic and Statistical Manual of Mental Disorders, Fifth Edition (DSM-5)[2]. Participants were recruited between March 2021 and March 2023 and assessed at the Central Institute of Mental Health, Mannheim, Germany. The EMA sample included n = 83 participants (53 men, 30 women; mean [SD] age, 37.20 [12.42] years; range, 17-62 years). General exclusion criteria included significant neurological conditions, substance use disorders other than alcohol, nicotine, or cannabis, head trauma, pregnancy, contraindications for magnetic resonance imaging, and a diagnosis of schizophrenia spectrum or bipolar disorders. The MRI sample included n = 80 participants after excluding 3 because of missing imaging data (n = 1) or insufficient image quality for CAT12 preprocessing (n = 2). The Institutional Review Board II of Heidelberg University, Germany, approved the study. All participants provided informed consent and received monetary compensation.

#### 1.2. Assessment procedures and measurements

##### Socio-demographic and psychological inventory

Clinical and psychological assessments included basic sociodemographic information (age, sex, education, and income) and a battery of established psychosocial self-report instruments, with validated German versions used. Education was self-reported as highest general schooling qualification and coded from 1 to 9: 1 = still in general full-time school; 2 = still in vocational-oriented school; 3 = left school without lower secondary qualification; 4 = lower secondary qualification (Hauptschule); 5 = intermediate secondary qualification (Realschule); 6 = polytechnic upper secondary qualification; 7 = university of applied sciences entrance qualification; 8 = general or subject-linked university entrance qualification (Abitur); 9 = other qualification. Monthly net personal income was self-reported and coded from 1 to 10: 1 = less than €500; 2 = €500 to €1,000; 3 = €1,000 to €1,500; 4 = €1,500 to €2,000; 5 = €2,000 to €2,500; 6 = €2,500 to €3,500; 7 = €3,500 to €4,000; 8 = €4,000 to €4,500; 9 = €4,500 to €5,000; 10 = more than €5,000. Alcohol-related baseline measures comprised current (past 12 months) and lifetime DSM-5 alcohol use disorder (AUD) criteria, assessed using the Structured Clinical Interview for DSM-5 (SCID-5)[3] and the Alcohol Use Disorders Identification Test (AUDIT)[4]. Perceived social support was assessed using the Oslo Social Support Scale (OSSS-3)[5], and social network structure was assessed using the Social Network Index (SNI)[6], yielding network size (SNI number) and network complexity (SNI complexity). Descriptive statistics and exploratory Spearman correlations with the Social Drinking Risk (SDR) index (individual estimate for surrounding drinkers derived from the multilevel model) are reported in Table S2, with false discovery rate (FDR) adjustment applied across the 10 correlations using the Benjamini-Hochberg procedure.

##### Ecological momentary assessments

###### *E-diary platform*

EMA was administered using Android smartphones and MovisensXS (XS-DFG v1.6; Movisens GmbH). Prompts were scheduled between 08:00 and 22:00 throughout the 6-week assessment period. Before monitoring, participants received standardized training on the smartphone and EMA procedures. Participants contributed 12,774 EMA observations (median [IQR], 163 [108-196] per participant). The GLMM included 5,221 completed non-solitary observations (median [IQR], 51 [28-87] per participant).

###### *Prompt schedule*

During weeks 1 to 3, participants received 8 prompts/day: 1 morning prompt randomized between 08:00 and 09:00 and 7 daytime prompts randomized between 09:00 and 22:00, spaced at least 1 hour apart. During weeks 4 to 6, participants received 4 prompts/day: 1 morning prompt randomized between 08:00 and 09:00 and 3 daytime prompts randomized between 09:00 and 22:00, spaced at least 2 hours apart. All morning and time-randomized daytime prompts were included as regular observations.

###### *E-diary measures*

The EMA items were administered in German; English translations are provided below. Three items were relevant to the present analysis:

1. Are you alone at the moment? (yes/no)

2. If not alone: How many individuals in your surrounding are drinking alcohol at the moment? (none, a few, about half, majority, all; coded 1-5)
3. Are you drinking alcohol at the moment? (yes/no)

#### *Multilevel modeling of e-diary assessments*

To test whether exposure to surrounding drinkers was associated with momentary drinking, we fitted a generalized linear mixed model with a logit link (PROC GLIMMIX) using SAS (version 9.4; SAS Institute Inc., Cary, NC, USA). The binary outcome was momentary drinking at observation i for participant j. The primary level-1 predictor was exposure to surrounding drinkers coded from 1 (none) to 5 (all) and modeled as a linear predictor. Time of day was included as a level-1 covariate; age and sex were included as level-2 covariates. The model included participant-specific random intercepts and random slopes for surrounding-drinkers exposure.

*Equation S1:*

$$logit[P(Y(\boldsymbol{Drinking\ at\ the\ moment})_{ij} = 1)] = \beta_{00} + \boldsymbol{\beta_{10}}(\mathbf{Surrounding\ Drinkers_{ij}}) + \beta_{01}(Sex_j) + \beta_{02}(Age_j) + \beta_{20}(Time\ of\ Day_{ij}) + u_{0j} + \boldsymbol{u_{1j}}(\boldsymbol{Surrounding\ Drinkers_{ij}})$$

where $Y_{ij}$ is drinking at the moment, $\beta_{10}$ is the fixed effect of surrounding drinkers, $u_{0j}$ is the participant-specific random intercept, and $u_{1j}$ is the random slope for surrounding drinkers. Participant-specific slope estimates (fixed effect plus random slope) for surrounding-drinkers exposure were extracted from this model and used as the SDR index in the neuroimaging analyses. The model results are presented in Table S1.

For Figure 1B, predicted probabilities at different surrounding-drinking values (1–5) were calculated for each participant using that participant's sex and age and a fixed time of day (18:00). Gray curves retained participant-specific random intercepts and slopes; the black curve used fixed effects only, and was averaged across participants at each exposure level. Corresponding linear predictors were plotted on the log-odds scale without inverse-logit transformation (Figure S1).

### Neuroimaging

T1-weighted structural images were acquired on a 3T Siemens MAGNETOM Prisma_fit (Siemens Healthineers, Erlangen, Germany) at the Central Institute of Mental Health (CIMH), Mannheim, using the 3D MPRAGE sequence (t1_mprage_sag_p2_iso; acquisition time 4:42 min). The imaging parameters were as follows: TR = 2000 ms, TE = 3.03 ms, TI = 900 ms, flip angle = 9°, voxel size = 1.0 × 1.0 × 1.0 mm, 192 slices, sagittal orientation, FOV = 256 mm, matrix = 256 × 256, and GRAPPA acceleration factor = 2. Structural images were preprocessed in MATLAB using SPM12 and the Computational Anatomy Toolbox. Processing used default CAT12 settings, including segmentation with the SPM12 tissue probability maps (TPM.nii), Dartel spatial normalization to the 1.5-mm IXI555_MNI152 template, MNI affine regularization, and SANLM denoising with MRF refinement (0.05). Internal CAT12 weighting parameters were LASstr = 0.50, GCUTstr = 0.50, and CLEANUPstr = 0.50. Modulated, spatially normalized gray matter maps were smoothed with an 8-mm full-width at half-maximum Gaussian kernel in SPM12. CAT12 report files were generated for each participant and used for quality control and extraction of global tissue volumes. The neuroimaging analyses included n = 80 participants after excluding 3 because of missing imaging data (n = 1) or insufficient image quality for CAT12 preprocessing (n = 2). Voxel-based morphometry (VBM)[7] was conducted in CAT12 within SPM12 (https://neuro-jena.github.io/cat/; www.fil.ion.ucl.ac.uk/spm). Smoothed, modulated gray matter images were entered into a voxel-wise multiple regression model across the brain (implicit gray matter mask). Subject-level covariates were SDR, total gray matter volume, age, and sex. The primary contrast tested the negative association of SDR with GMV. Reported peaks were evaluated with family-wise error small-volume correction (FWE-SVC) within a bilateral pgACC mask defined a priori (see below). The peak negative association was in the pgACC (MNI, [2, 48, 10]; T = 5.40; P < .001, FWE-SVC). Subject-level pgACC GMV was then extracted at this peak from each participant's modulated gray matter image using spm_summarise (SPM12). This peak-voxel pgACC GMV was used for Fig. 2A and as a subject-level covariate in the structural covariance model. For the structural covariance analysis, following established interregional covariance approaches[8], a second voxel-wise model was specified on the same gray matter images. Covariates were peak-voxel pgACC GMV, SDR, their product (peak-voxel pgACC GMV × SDR), total gray matter volume, age, and sex. The primary contrast tested the pgACC × SDR interaction term. Maps were estimated voxel-wise across the brain; reported peaks were evaluated with FWE-SVC within the a priori VS mask. The interaction was significant at the VS peak (MNI, [8, 15, -9]; T = 3.15; P = .032, FWE-SVC), consistent with weaker pgACC–VS structural covariance among participants with higher SDR. For Figure 2C, peak-voxel VS GMV was extracted at this coordinate with spm_summarise and plotted against peak-voxel pgACC GMV; groups were split by the median SDR for illustration only and were not used for inference. A priori small volumes for FWE-SVC were bilateral pgACC (VBM) and VS (structural covariance), delineated with the Automated Anatomical Labeling 3 (AAL3) atlas[9]. The VS mask was further refined using a gray matter probability threshold greater than 80% based on the tissue probability map. Sagittal brain maps in Figure 2 were rendered in MRIcroGL and overlaid on the standard MNI152 T1-weighted template distributed with MRIcroGL (mni152.nii.gz), at P < .05, uncorrected, for display; statistical inference used FWE-SVC as above.

### 2. Supplementary Tables

#### 2.1. Table S1: Association of Exposure to Surrounding Drinkers With Momentary Drinking in the Generalized Linear Mixed Model

| | **Estimate** | **Standard error** | **OR** | **95% CI** | **DF / 95% bounds** | **Test statistic** | **p-value** |
|---|---|---|---|---|---|---|---|
| **Fixed effects** | | | | | | | |
| Intercept | −10.332 | 0.6073 | <.001 | <.001–<.001 | 80 | −17.01 | <.0001 |
| Exposure to surrounding drinkers | 1.4419 | 0.08114 | 4.23 | 3.60–4.97 | 75 | 17.77 | <.0001 |
| Sex (female) | −0.3615 | 0.2871 | 0.70 | 0.40–1.22 | 5061 | −1.26 | 0.2081 |
| Age, y | 0.02612 | 0.0116 | 1.03 | 1.00–1.05 | 5061 | 2.25 | 0.0244 |
| Time of day, h | 0.2868 | 0.01952 | 1.33 | 1.28–1.38 | 5061 | 14.69 | <.0001 |
| **Random Effects** | | | | | | | |
| Intercept | 0.7332 | 0.2395 | — | — | 0.4227 to 1.5737 | z = 3.06 | 0.0011 |
| Exposure to Surrounding drinkers | 0.1415 | 0.05544 | — | — | 0.07438 to 0.3668 | z = 2.55 | 0.0054 |

CI = confidence interval; DF = degrees of freedom; EMA = ecological momentary assessment; GLMM = generalized linear mixed model; h = hours; OR = odds ratio; y = years. Fixed-effect estimates are expressed on the log-odds scale. Odds ratios and 95% CIs were obtained by exponentiating the estimates and their SAS-reported 95% confidence limits. The odds ratio for surrounding-drinkers exposure corresponds to a one-point increase. Fixed-effect P-values are reported 2-sided. Random intercepts and slopes are modeled as independent using a variance-components covariance structure.

#### 2.2. Table S2: *Sample statistics and correlations with the SDR index.*

| | **EMA sample** | | | **Correlation with SDR** | | |
|---|---|---|---|---|---|---|
| | M ± SD / count | min/max | n | *Spearman's ρ* | *p*-value | *FDR-adjusted p-value* |
| ***Demographics*** | | | | | | |
| Age (years) | 37.20 ± 12.42 | 17/62 | 83 | 0.059 | 0.598 | 0.748 |
| Sex (53 male / 30 female) | | | 83 | −0.132 | 0.235 | 0.584 |
| Education | 6.71 ± 1.83 | 1/9 | 82 | −0.022 | 0.848 | 0.848 |
| Monthly net income | 4.68 ± 2.56 | 1/10 | 82 | 0.090 | 0.422 | 0.603 |
| ***Alcohol use*** | | | | | | |
| DSM-5 AUD criteria, current 12 months | 4.02 ± 1.59 | 2/8 | 83 | 0.117 | 0.292 | 0.584 |
| DSM-5 AUD criteria, lifetime | 4.17 ± 2.29 | 0/11 | 83 | 0.266 | 0.015 | 0.110 |
| AUDIT total | 14.38 ± 5.84 | 3/29 | 82 | −0.038 | 0.734 | 0.816 |

| | EMA sample | | | Correlation with SDR | | |
|---|---|---|---|---|---|---|
| ***Psychosocial*** | | | | | | |
| OSSS-3<br>social support | 11.24 ± 2.14 | 6/14 | 82 | −0.102 | 0.360 | 0.600 |
| SNI network size | 17.10 ± 11.72 | 4/60 | 82 | 0.231 | 0.037 | 0.123 |
| SNI network complexity | 5.96 ± 2.06 | 2/10 | 82 | 0.253 | 0.022 | 0.110 |

*AUD = alcohol use disorder; AUDIT = Alcohol Use Disorders Identification Test; OSSS-3 = Oslo Social Support Scale; SNI = Social Network Index; SDR = social drinking risk; SD = standard deviation; FDR = false discovery rate. Education and monthly net income were self-reported variables coded as described in section 1.2. FDR-adjusted p-values were computed using the Benjamini-Hochberg procedure across the 10 correlations in this table. Missing values resulted from participants opting not to disclose certain information or from non-response to individual items.*

### 3. Supplementary Figures

#### 3.1. Figure S1: *Predicted log odds of momentary drinking by exposure to surrounding drinkers.*

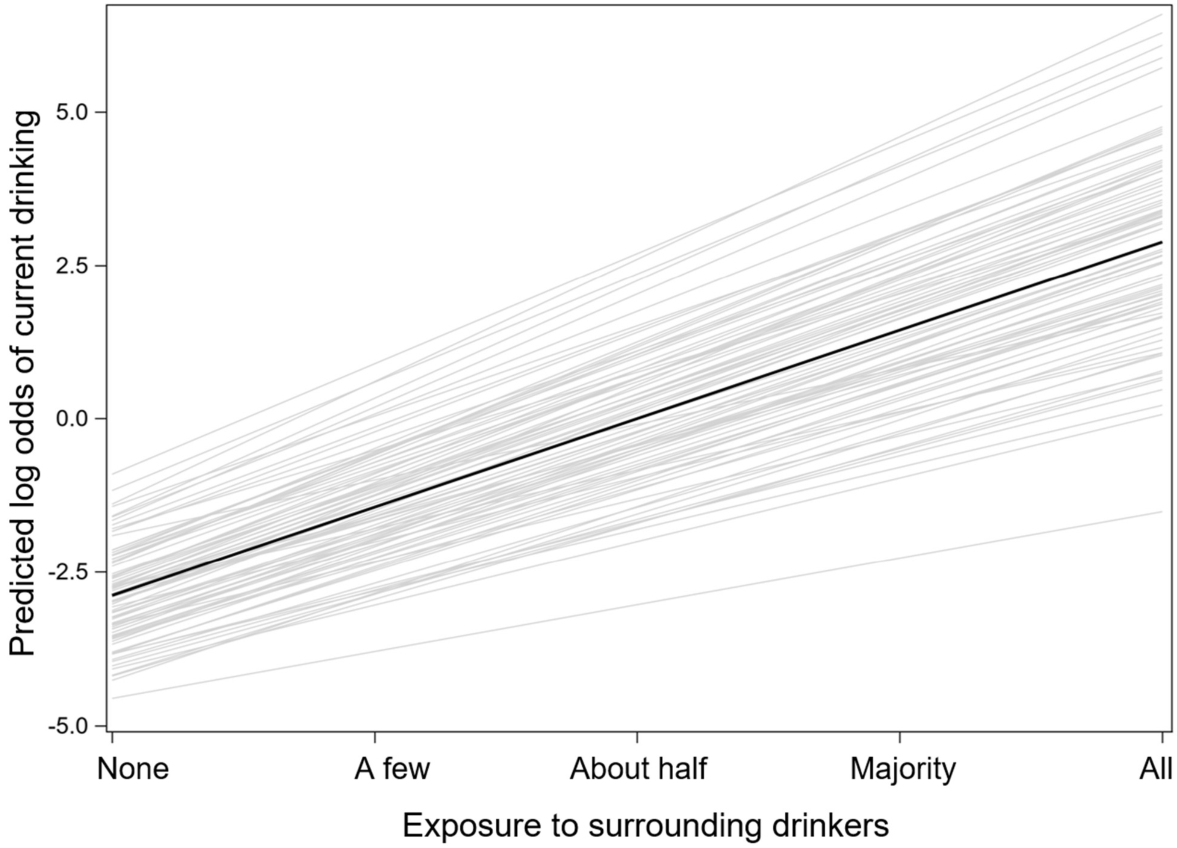

Participant-specific predictions on the model's log-odds scale. For each participant, predictions were evaluated at that participant's sex and age and a fixed time of day (18:00). Gray lines include participant-specific random intercepts and slopes; the black line shows the fixed-effects prediction averaged across participants at each exposure level.

### Supplementary References

1 Heinz, A., Kiefer, F., Smolka, M. N., Endrass, T., Beste, C., Beck, A. *et al.* Addiction Research Consortium: Losing and regaining control over drug intake (ReCoDe)-From trajectories to mechanisms and interventions. *Addict Biol* **25**, e12866 (2020). https://doi.org/10.1111/adb.12866

2 American Psychiatric Association. *Diagnostic and statistical manual of mental disorders : (DSM-5®)*. Fifth edition. edn, (American Psychiatric Association, 2013).

3 First, M., Williams, J., Karg, R. & Spitzer, R. *Structured Clinical Interview for DSM-5 Disorders, Clinician Version (SCID-5-CV)*. (American Psychiatric Association Publishing, 2016).

4 Saunders, J. B., Aasland, O. G., Babor, T. F., de la Fuente, J. R. & Grant, M. Development of the Alcohol Use Disorders Identification Test (AUDIT): WHO Collaborative Project on Early Detection of Persons with Harmful Alcohol Consumption--II. *Addiction* **88**, 791-804 (1993). https://doi.org/10.1111/j.1360-0443.1993.tb02093.x

5 Kocalevent, R. D., Berg, L., Beutel, M. E., Hinz, A., Zenger, M., Harter, M. *et al.* Social support in the general population: standardization of the Oslo social support scale (OSSS-3). *BMC Psychol* **6**, 31 (2018). https://doi.org/10.1186/s40359-018-0249-9

6 Cohen, S., Doyle, W. J., Skoner, D. P., Rabin, B. S. & Gwaltney, J. M., Jr. Social ties and susceptibility to the common cold. *JAMA* **277**, 1940-1944 (1997).

7 Ashburner, J. & Friston, K. J. Voxel-based morphometry--the methods. *Neuroimage* **11**, 805-821 (2000). https://doi.org/10.1006/nimg.2000.0582

8 Mechelli, A., Friston, K. J., Frackowiak, R. S. & Price, C. J. Structural covariance in the human cortex. *J Neurosci* **25**, 8303-8310 (2005). https://doi.org/10.1523/JNEUROSCI.0357-05.2005

9 Rolls, E. T., Huang, C. C., Lin, C. P., Feng, J. & Joliot, M. Automated anatomical labelling atlas 3. *Neuroimage* **206**, 116189 (2020). https://doi.org/10.1016/j.neuroimage.2019.116189
